# Sequence Alignment on Directed Graphs

**DOI:** 10.1101/124941

**Authors:** Kavya Vaddadi, Naveen Sivadasan, Kshitij Tayal, Rajgopal Srinivasan

**Author notes:** Email: {, }.

## Abstract

Genomic variations in a reference collection are naturally represented as genome variation graphs. Such graphs encode common subsequences as vertices and the variations are captured using additional vertices and directed edges. The resulting graphs are directed graphs possibly with cycles. Existing algorithms for aligning sequences on such graphs make use of partial order alignment (POA) techniques that work on directed acyclic graphs (DAG). For this, acyclic extensions of the input graphs are first constructed through expensive loop unrolling steps (DAGification). Also, such graph extensions could have considerable blow up in their size and in the worst case the blow up factor is proportional to the input sequence length. We provide a novel alignment algorithm V-ALIGN that aligns the input sequence directly on the input graph while avoiding such expensive DAGification steps. V-ALIGN is based on a novel dynamic programming formulation that allows gapped alignment directly on the input graph. It supports affine and linear gaps. We also propose refinements to V-ALIGN for better performance in practice. In this, the time to fill the DP table has linear dependence on the sizes of the sequence, the graph and its feedback vertex set. We perform experiments to compare against the POA based alignment. For aligning short sequences, standard approaches restrict the expensive gapped alignment to small filtered subgraphs having high ‘similarity’ to the input sequence. In such cases, the performance of V-ALIGN for gapped alignment on the filtered subgraph depends on the subgraph sizes.

## I. INTRODUCTION

Most state-of-the-art high throughput genome studies rely heavily on high quality reference genome [1]. Single reference sequence however has limited capability in representing significant genomic diversities and it suffers from reference allele bias during interpretations [2], [3], [4]. The number of sequenced genomes is ever increasing and this is driving a paradigm shift in genome analysis from single reference sequence based to pangenome reference based [2], [3], [4].

Representing the genomic variations using graph data structures have attracted considerable interest recently [2], [3], [4], [5], [6]. Various graph data structures have been studied in the literature for pangenome representation with subtle distinctions [3]. These include De Bruijn graphs [7], [8], A-Bruijn graphs [9], Enredo graphs [10], Cactus graphs [5], [11], Population Reference graphs [6], String graphs [12], and Variation graphs [2]. The broad idea behind these representations is to effectively encode complex genomic variations such as insertions, deletions, duplications, transpositions, reversals, rearrangements etc., as alternative paths in the graph using additional edges and vertices. The resultant graphs are directed and may contain cycles. In variation graphs [2], the common subsequences are encoded as labeled vertices and variations are represented using additional vertices and directed edges. Such representations have shown promise in improved read mapping and variant calling performance [4]. Graph based reference has necessitated the development of graph based computational pipelines for genome analyses [3], [2], [4].

Sequence alignment is a fundamental problem in genomics. In this paper, we consider the alignment of a sequence to a pangenome reference which is encoded as a graph. In the graph, common subsequences are represented as vertices which are labeled by the subsequences they encode. The variations are captured using additional labeled vertices and directed edges. Representing variations such as duplications and highly varying copy numbers in these graphs could introduce directed cycles. The sequences in the pangenome are present as directed paths (not necessarily simple) in the graph. Our goal is to compute an alignment of the input sequence to a path in the graph having maximum alignment score among all paths in the graph. We consider gapped alignments where the gaps could be affine, linear or constant. The formal problem definitions are given in Section II.

Algorithm for aligning a new sequence to a multiple sequence alignment (MSA) encoded as graph was given in [13]. MSA is encoded as a Partial Order Alignment graph (POA) and the alignment algorithm aligns the new sequence to the POA [13]. The POA based alignment algorithm of [13] is an extension of the traditional dynamic programming algorithms for sequences [14], [15] to handle partial orders. In POA graphs, sequences are represented as paths and each of the vertices can have multiple incoming and outgoing edges. POA graphs are Directed Acyclic Graphs (DAG). Gapped alignment of an *n* length sequence to a POA graph on *E* edges takes *O*(*mE*) time [13].

The POA graphs share resemblance to reference graphs in the sense that the variations are encoded as alternative paths using additional vertices and edges. In [4], POA based alignment was used for aligning sequences to reference graphs. POA graphs however acyclic. Capturing complex structural variations such as inversions, duplications or copy numbers with high variability could result in back edges and consequently directed cycles in the graphs. Genome graphs such as variation graphs [2] that attempts to capture such variations are hence not necessarily acyclic. The POA based alignment algorithm cannot be used directly on such graphs. To handle such graphs, acyclic extensions of the input graphs are constructed through expensive loop unrolling (*DAGification*) steps [4] and the alignment is then performed on the acyclic extensions. A *k*-length DAGification of graph *G* aims to compute a DAG *G*’ such that all paths (not necessarily simple) of length *k* or less in *G* are present in *G*’ and vice versa. For aligning an *m* length sequence, the value of *k* has to be *m* or more. Such graph extensions can however have considerable increase in their size. The edge and vertex blow up factor in the worst case is proportional to the input sequence length. Prohibitively large size of the DAGified graph results in increased preprocessing and alignment time and thereby affects the overall alignment performance.

Performance of short sequence alignment and read mapping to large reference graphs is improved by a filtering phase. In this phase, candidate subgraphs of the reference graph with potentially large alignment score with the input sequence are identified [2], [4]. The final alignment is then computed by performing alignment on each of these candidate subgraphs and choosing the best. In this case, the DAGification is restricted to the candidate subgraphs.

### A. Our Contribution

In this paper, we provide a novel alignment algorithm V-ALIGN that aligns the input sequence directly on the input graph while avoiding expensive DAGification preprocessing. It computes an alignment of the input sequence to a path in the graph having maximum alignment score among all paths in the graph. V-ALIGN is based on a novel dynamic programming formulation that allows gapped alignment with affine, linear or constant gaps directly on the input graph. We also propose refinements to V-ALIGN for better performance in practice. In this, the time to fill the DP table has linear dependence on the sizes of the sequence, the graph and its *feedback vertex set*. A feedback vertex set of a graph is a subset of its vertices whose removal makes the graph acyclic. The runtime of this algorithm matches that of the POA based alignment when the reference graph is acyclic. V-ALIGN performs one time preprocessing of the graph to compute pairwise edge distances between the vertices and to compute a feedback vertex set. When the alignment is restricted to a filtered set of subgraphs, which is done for improved efficiency, the V-ALIGN can be used for aligning to these candidate subgraphs. In this case, its performance depends on the subgraph sizes. We also provide a theoretical result on the complexity of the DAGification preprocessing which is required by the POA based alignment. We show that 2-length DAGification of a graph *G* where the resultant DAG has minimum number of vertices is NP-complete. We also conduct empirical studies on the DAGification overhead. For this, we use a *dfs* based DAGification algorithm and measure the blow up in the vertices and edges of the resultant graphs for different types of input graphs.

## II. PRELIMINARIES

### A. Notations

Let *G* = (*V,E, γ*) be a connected directed graph with vertices *V*, edges *E* and vertex labels given by *γ*(*v*). Edges in *E* are represented as ordered pairs from *V* × *V*. Let Σ^+^ denote the set of all sequences of one or more elements from an alphabet Σ. For nucleotide sequences, Σ is the set of possible nucleotides. For a vertex *v* ∈ *V*, its label *γ*(*v*) ∈ Σ^+^. A directed path *p* in *G* of length *r* vertices is denoted by the ordered sequence (*u*_1_,…,*u*_*r*+1_), where *u_i_* ∈ *V* and (*u_i_, u*_*i*+1_) ∈ *E*. We only consider paths with length greater than 0. We say that the path *p* starts at *u*_1_ and ends at *u_r_*. Let *P* (*v*) denote the set of all directed paths in *G* that end at vertex *v*. Clearly the cardinality of *P* (*v*) could be infinity if there are directed cycles in *G*. For an ordered sequence *x* = (*x*_1_, *x*_2_,…,*x_m_*), let |*x*| denote the length of sequence *x*, which is *m* here. For a directed path *p* = (*u*_1_,…,*u_t_*) in *G* (not necessarily simple), we call the sequence obtained by concatenating *γ*(*u*_1_),…,*γ*(*u_t_*) in the same order as the label of the path *p* and is denoted by *γ*(*p*). Hence 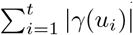. For any contiguous subsequence *y* of *γ*(*p*), we say that *G* contains the label sequence *y*. For a vertex *v* ∈ *V*, let *I*(*v*) called the *in-neighbors* of *v* denote the set of all vertices that have directed edges to *v*. For a set *E*, we also use *E* to denote its cardinality in place of |*E*| inside asymptotic notations for better readability.

### B. Alignment Problems

We consider gapped alignment of a sequence *x* ∈ Σ^+^ to *G*, where the goal is to compute an alignment of *x* to *γ*(*p*) for some path *p* in *G*, that achieves the maximum alignment score among all paths in *G*. Since *G* is a directed graph possibly with cycles, the standard global alignment and local alignment between sequences translate to the following two variants:

- End to end alignment of *x* to a contiguous sub sequence *y* of *γ*(*p*), without penalizing the unmatched suffix and prefix in *γ*(*p*). The maximum alignment score is denoted as *g*(*x, γ*(*p*)).
- Local alignment of *x* and *γ*(*p*). The maximum alignment score is denoted as *l*(*x, γ*(*p*)).

Consequently, we define 
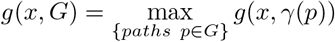
and 
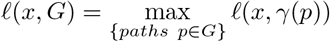

## III. ALIGNMENT ALGORITHMS

For the ease of exposition, we assume that the vertex labels are length one sequences from Σ^+^. That is, the label of any vertex in *G* is an element from Σ. We will discuss later how the algorithm can be easily modified to handle the general case. In the following, we define a dynamic programming formulation that would allow us to find optimal alignments.

For vertices *u* and *v* in *G*, let *δ*(*u, v*) denote the minimum number of edges on any directed path from *u* to *v* in *G*. That is, *δ*(*u,v*) is the shortest edge distance from *u* to *v*. Clearly *δ*(*u,u*) = 0. If there is no directed path from *u* to *v* then *δ*(*u,v*) = +*∞*. For *a, b* ∈ Σ, let *s*(*a,b*) denote the substitution score between *a* and *b*. Let Δ(*k*) denote the penalty for a *k* length gap, such as affine, linear or constant gap. Δ(0) = 0 by definition.

Let the input sequence *x* = (*x*_1_,…,*x_m_*) of length *m*. We assume an arbitrary linear ordering of the vertices in *V*. Let *M* denote the scoring matrix of size |*V*| × (*m* + 1) where *M* (*w, j*) is the entry for *w* ∈ *V* and *j* ∈ [0, *m*]. We use the following recurrence relation on *M* (*w, j*) for all *j* ∈ [1, *m*] and *w* ∈ *V*:

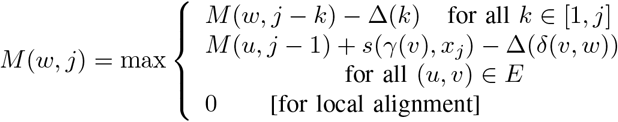

The entry *M* (*w, j*) stores the maximum score for aligning the subsequence (*x*_1_,…,*x_j_*) to any path ending at vertex *w* in *G*. The first term of the above max expression corresponds to an alignment having *k* gaps in the end due to the deletion of the last *k* elements of (*x*_1_,…,*x_j_*). The second term corresponds to aligning *x_j_* to an intermediate vertex *v* in the path followed by gaps due to the deletion of the remaining path, which is no more than *δ*(*v,w*) for an optimal alignment.

There could be vertices in *G* with no incoming edges (zero in-degree). In order to handle such vertices, we always include a dummy vertex *θ* in the vertex set *V* and add directed edges from *θ* to each vertex in *G* with zero in-degree. Matrix *M* is initialized as *M* (*w*,0) = 0 for each *w* ∈ *V* and *M* (*θ, j*) = 0 for all *j* ∈ [0, *m*] for local alignment. For the score function *g*(), the 0 term is absent from the max expression in the above recurrence and *M* is initialized as *M* (*w*,0) = 0 and *M* (*θ, j*) = −Δ(*j*) for *j* ∈ [1, *m*]. Computing alignment score *ℓ*(*x,G*) and an alignment path from *M* are done in the usual manner. For *g*(), the alignment score is the largest *M* (*v,m*) entry.

The computational efficiency can be improved further using standard techniques [15] (for affine, linear or constant gaps) by defining an auxiliary matrix *Q* where 
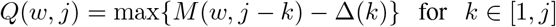
and replacing the first term in the max expression above with *Q*(*w, j*). Value of *Q*(*w, j*) can be updated in *O*(1) time because of the recurrence *Q*(*w, j*) = max{*M*(*w, j* − 1) − Δ(1), *Q*(*w, j* − 1) − *t*}, where *t* is the gap extension cost. *Q*(*w*, 0) = −∞ for all *w*. The time complexity for filling *M* is *O*(*mV E*). Computing *δ*(*u,v*) is a one time preprocessing which can be done in *O*(*V E*) time.

### A. Improved V-ALIGN Algorithm

We now provide a modified dynamic programming formulation to compute *M* which can achieve better run time performance in practice. Consider some linear ordering of the vertices in *V*. We can assume that the dummy vertex *θ* is the first vertex in the ordering. A vertex *v* is called *in-order* with respect to this given ordering if all vertices in *I*(*v*) (in-neighbors of *v*) lie to the left of *v* in this ordering. Let *V*′ ⊆ *V* denote the set of all vertices that are not in-order. We note that if *G* is acyclic (DAG) then the topological sorting gives an ordering where *V*′ = ∅. In directed graphs with cycles, |*V*′| > 0. If *G* can be made acyclic (DAG) by removing at most *α* edges then clearly |*V*′| ≤ *α*. This is because, introducing all the deleted *α* edges to a topological sorted order of the DAG can make at most *α* vertices not in-order.

We assume that the matrix rows are permuted with respect to the linear ordering of *V*. The technique in [15] for sequences can be extended to handle our case as follows. The earlier recurrence for *M* (*w, j*) can be rewritten as follows. For *j* ∈ [1, *m*] and for all *w* except *θ*,

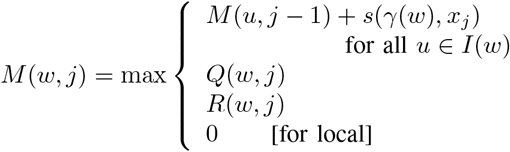
where 
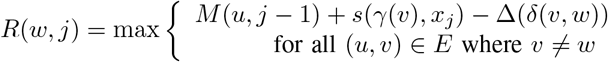

As earlier, *Q* can be computed efficiently using the recurrence

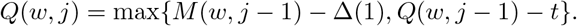

We recall that the matrix rows are permuted with respect to the linear ordering of *V*. This ensures that while computing the matrix entry (*w, j*) for some vertex *w* ∈ *V* − *V*′, the values of the entries (*u,j*) for all *u* ∈ *I*(*v*) (the in-neighbors of *v*) are already available. Hence, if *w* ∈ *V* − *V*′ then *R*(*w, j*) can be computed efficiently using the recurrence

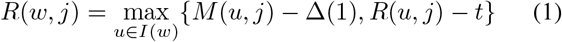

Value of *t* in the *Q*(*w, j*) and *R*(*w, j*) expressions above is the gap extension cost. Matrices *M* and *Q* are initialized as described earlier and *R*(*θ, j*) = −∞ for *j* ∈ [1,*m*].

While filling any column *j* of matrix *R*, the *R*(*w, j*) entries for *w* ∈ *V*′ are filled first using its original definition and the remaining entries for vertices in *V* − *V*′ are filled using (1).

### B. Time Complexity

We note that the time for updating one column of *M* is the sum total of all in degrees of vertices in *V*, which is *O*(*E*). Time to update one column of *Q* is *O*(*V*). Time for updating one column of *R* is the sum of the time taken for all *v* ∈ *V*′ and the time taken for all *v* ∈ *V* − *V*′. The first component is *O*(*V*′ *E*) and the second component is the sum of in degrees, which is again *O*(*E*). Hence the total time for filling *R* is *O*(*mE*(*V*′ + 1)). If *V*′ = ∅, which is the case when *G* is a DAG, then the run time matches [13]. We note that in general, there are graphs where the number of vertices in *V*′ can be Ω(*V*) for any ordering. The run time for such graphs is *O*(*mV E*) which matches the run time of our basic algorithm.

If *f* is the minimum number of vertices whose removal makes *G* acyclic, which is called the *minimum feedback vertex set* (MFVS), then the time taken in *O*(*m*(*f* + 1)*E*). This is because, we can use the vertex ordering obtained by topological sorting of *V* minus MFVS vertices and then place the MFVS vertices in the beginning of this ordering. Depending on *f*, the runtime could be smaller than *O*(*mV E*) time for our basic algorithm. Though MFVS problem is NP-complete, approximation algorithms, parameterized algorithms and efficient exact algorithms for special graph classes are known [16], [17], [18], [19], [20]. Any ordering with small *V t* set can lead to improved performance in our case. We also remark that if *G* is a simple directed path then the alignment problem reduces to the alignment of two sequences, of lengths |*V*| and *m* respectively, and the time taken in this case is *O*(*mE*) = *O*(*mV*) = *O*(*mn*) where |*V*| = *n*.

### C. Vertex Labels

In the previous section we assumed that the vertex labels are elements from Σ. We extend our algorithm in a straightforward manner to handle vertex labels from Σ^+^. Consider the directed graph *G_a_* = (*V_a_, E_a_, γ*) derived from *G* as follows. For each vertex *v* ∈ *V*, with label *γ*(*v*) = (*x*_1_,…,*x_r_*), include a chain of *r* vertices *v*_1_,…,*v_r_* in *G_a_* with directed edges from *v_i_* to *v*_*i*+1_. Label of *v_i_* is given by *γ*(*v_i_*) = *x_i_*. Clearly |*V_a_*| = Σ_*v*∈*V*_ |*γ*(*v*)|. For each directed edge (*u, v*) ∈ *E*, we include a directed edge (*u*_|*γ*(*v*)|_, *v*_1_) to *E_a_*. Hence |*E_a_*| = |*E*| + Σ_*v*∈*V*_ (|*γ*(*v*)| − 1). The linear ordering of *V* can be extended easily to obtain a linear ordering of *V_a_* by replacing each vertex *v* in the linear ordering by the corresponding chain of vertices *v*_1_,…,*v*_|*γ*(*v*)|_ in the new ordering. For any vertex *v* ∈ *V*, clearly the corresponding vertices *v*_2_,…,*v*_|*γ*(*v*)|_ ∈ *V_a_* are in-order vertices and *v*_1_ is in-order vertex if and only if *v* is in-order vertex. That is, for *G_a_*, the set of vertices not in-order is given by 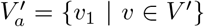. Hence 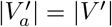. The dynamic programming matrix *M_a_* is of size |*V_a_*|× *m* and it is filled in the same manner.

The time for filling *M* is now *O*(*mE_a_*(*V*′ +1)) = *O*(*m*(*n*+ *E*)(*V*′ + 1)) where *n* = Σ_*v*∈*V*_ |*γ*(*v*)| is the sum total of the sizes of all vertex labels in *G*. For constant length vertex labels, the time is thus *O*(*mE*(*V*′ + 1)). If *G* is acyclic then the time is *O*(*mE*) for constant length vertex labels. We remark that if *G* is just a single vertex with a sequence label of length *n* and containing no edges, the alignment problem reduces to the standard sequence to sequence alignment. In this case, V-ALIGN takes *O*(*mn*) time.

We do not require the pre-computation of all-pair shortest edge distances in *G_a_*. Instead, we can compute vertex weighted all-pair shortest paths in *G* and use them to obtain *δ*(*u_i_, v_j_*) for any pair of vertices *u_i_, v_j_* in *G_a_*. Each vertex *v* in *G* assigned a weight *w*(*v*) equal to the length of its label. That is, *w*(*v*) = |*γ*(*v*)|. Weight of a path in *G* is the sum of the vertex weights in the corresponding vertex sequence. Now we compute *δ_w_* (*u, v*) for *u, v* in *G* which is the minimum weight of any path from *u* to *v*. We define *δ_w_* (*u, v*) = ∞ if there is no path from *u* to *v*. For *u* ≠ *v*, the shortest edge distance from *u_i_* to *v_j_* in *G_a_* is now given by *δ*(*u_i_, v_j_*) = *j*+*δ_w_* (*u, v*)−*i*−|*γ*(*v*)|. Clearly *δ*(*u_i_, u_j_*) = *j* − *i* where *j* ≥ *i*. Computing *δ_w_* for allpairs can be done using the standard Dijkstra’s algorithm in *O*(*VE* + *V*^2^ log *V*) time.

### D. Comparison with POA based alignment

The DAGification preprocessing in POA based alignment blows up the number of vertices and edges in the DAGified graph. The size of the DAGified graph depends on the topology of the input graph *G* and the input sequence length *m*. If *r* is the length of the shortest directed cycle in *G*, called the *girth* of *G*, then the DAGification will unroll this cycle Θ(*m*/*r*) times. There are graphs where this can result in a blow up of vertices and edges by a multiplicative factor Θ(*m*/*r*). Such graphs are discussed in the experiments section (Section V). The time complexity for POA based alignment is such cases is *O*(*mE*(*m*/*r* + 1)). In the worst case, this can be *O*(*m*^2^*E*), which grows quadratically with the input sequence length. This time complexity is excluding the time required for the DAGification.

### E. Aligning Short Sequences

Usual approaches for fast alignment of a short sequence *x* to a large target sequence *y* follows efficient filtering of regions (subsequences) in *y* having high ‘similarity’ with *x* and restricting expensive gapped alignment only to these regions. In the same manner, for aligning to a large graph *G*, alignment can be restricted to regions (subgraphs) of *G* having high ‘similarity’. Such subgraphs can have considerably lesser number of vertices and edges as compared to *G* and this can lead to faster alignment. Even though the shortest edge distances between vertex pairs (*u, v*) can be different (higher) in the subgraph, V-ALIGN can still use the *δ*(*u, v*) values computed on *G* for the subgraph alignment. From the algorithm description, it is clear that this can only result in a possibly better alignment of *x* to *G*. This is because, the shortest paths correspond to regions of deletions along the candidate paths in *G* while aligning with *x*. On the other hand, the vertex linear ordering restricted to a subgraph be better compared to the linear ordering of all vertices in *G*. Recomputing the ordering for subgraphs is a matter of choice based on performance considerations.

## IV. COMPLEXITY OF DAGIFICATION

We present results on the complexity of DAGification preprocessing used by POA based alignment techniques. We assume that the directed graph *G* = (*V, E, γ*) is connected and that the vertex labels are elements of the alphabet Σ. We first define a *k*-DAG of a directed graph *G* for *k* ≥ 1. We say that a directed acyclic graph *G*′ = (*V*′, *E*′, *γ*′) is a *k*-DAG of *G* when the following holds: *G* contains a label sequence *y* of length *k* or less if and only if *G*′ contains *y*. For a graph *G* = (*V, E, γ*), clearly *G*′ = (*V*, ∅, *γ*) is a 1-DAG of *G*. A *k*-DAG of *G* with *k* ≥ 2 may contain more vertices and edges than *G*. This is because, additional vertex copies with the same vertex label are included in *G*′ every time the same vertex is encountered during loop unrolling.

POA based sequence alignment first computes a *k*-DAG for the input graph *G* and then computes an optimal gapped alignment of the input sequence to some label sequence contained in the *k*-DAG. If the vertex labels in *G* are just the elements of the alphabet Σ then the gapped alignment of an *m* length sequence requires a *k*-DAG of *G* with *k* ≥ *m*. Aligning the *m* length sequence on a *k*-DAG *G*′ = (*V*′, *E*′, *γ*′) requires *O*(*m*|*E*′|) time using [13]. Hence the size of the *k*-DAG affects the alignment performance.

In the following we present a simple complexity result on computing *k*-DAG of a directed graph *G* = (*V, E, γ*) assuming that vertex labels in *G* are elements of Σ and the labels are distinct. For simplicity, we assume that Σ = *V* and *γ*(*v*) = *v*.

**Theorem 1.** *For directed graph G* = (*V, E, γ*), *computing a* 2-*DAG of G having minimum number of vertices is NP-complete*.

*Proof*. Clearly *G* is assumed to be not a DAG. Let *V_f_* = {*u*_1_,…,*u_f_*} be a feedback vertex set of *G*. Consider the graph *G*′ obtained by adding *f* new vertices {*v*_1_,…,*v_f_*} to *G* where the label of *v_i_* is *γ*(*u_i_*). Each directed edge (*w, u_i_*) for any *u_i_* ∈ *V_f_* is replaced in *G*′ by a new edge (*w, v_i_*). It is straightforward to verify that *G*′ is a 2-DAG of *G* with |*V*| + *f* vertices. Conversely, if *G*″ = (*V*″, *E*″, *γ*″) is a 2-DAG of *G*, then the set of vertex labels each of which is assigned to more than one vertex in *G*″ forms a feedback vertex set of *G*. Its cardinality at most |*V*″| − |*V*|. The result now follows from the NP-completeness of MFVS computation on directed graphs. □

Generating a *k*-DAG for an input graph *G* can nevertheless be done with a simple *k*-depth *dfs* traversal that explores *k* length paths in *G* [2]. The output DAG however need not have the minimum number of vertices or edges. Each strongly connected component of *G* can be DAGified separately. For now we assume *G* is strongly connected. A separate *k*-depth dfs starting from each vertex of *G* is performed. Each vertex of the DAG has an associated level. During the traversal from a vertex, its neighboring vertices in *G* and the connecting edges are added to the same level in the DAG unless the neighbor is already present at that level. Otherwise, an additional copy of the neighboring vertex is added to the next level and the connecting edge goes across the two levels. The traversal now proceeds from the newly added copy of the neighbor. Vertices are added to level 0 when encountered for the first time. Repeated invocations of an *r*-depth dfs from a vertex copy for the same *r* value is avoided by book keeping. Output of such a DAGification procedure is used in the experiments discussed in the next section.

## V. EXPERIMENTS

We conduct experiments to study the increase in the size of the input graph due to the *k*-DAGification preprocessing. This preprocessing is required by the POA based techniques whereas our algorithm avoids this expensive computation. The candidate set of graphs *C* used in our experiments consists of two classes of synthetic graphs. The first class consists of complete graphs *K_n_* with *n* ranging from 1 to 5. A *K*_3_ is shown in Figure 2. Girth of *K_n_* is 1. The second class of graphs have larger girth values. Figure 1 shows the graphs in this class with girth value *r* ranging from 2 to 4. For each of these graphs, its multiple copies are connected to form a chain graph with the same girth value. These chain graphs are also included in the second class. Figure 2 shows a chain graph obtained from the girth 3 graph. In this graph, vertices 5, 9 and 13 are copies of vertex 1.

**Fig. 1:**
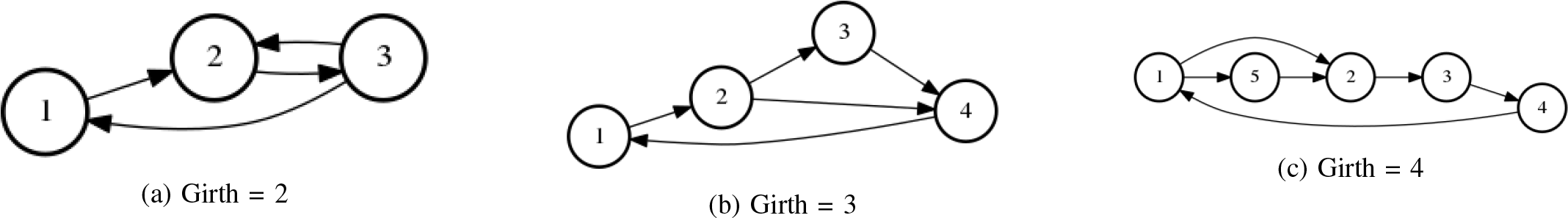
Graphs with varying girth

**Fig. 2:**
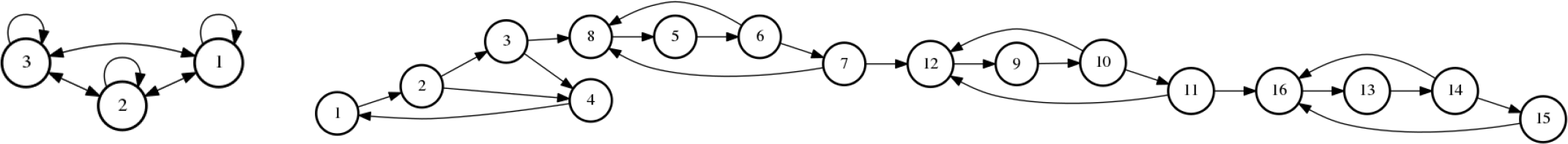
The left side graph is *K*_3_. The right side graph is a chain of 4 copies of the girth 3 graph (Fig. 1b).

Figure 3 shows the increase in the vertices and edges in the *k*-DAGification output for all the candidate graphs in 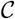. The value of *k* depends on the length of the sequence to be aligned and *k* should be greater than the sequence length to allow for gaps in the alignment. We consider *k* values in the range [10, 50] and in the range [100, 1000].

**Fig. 3:**
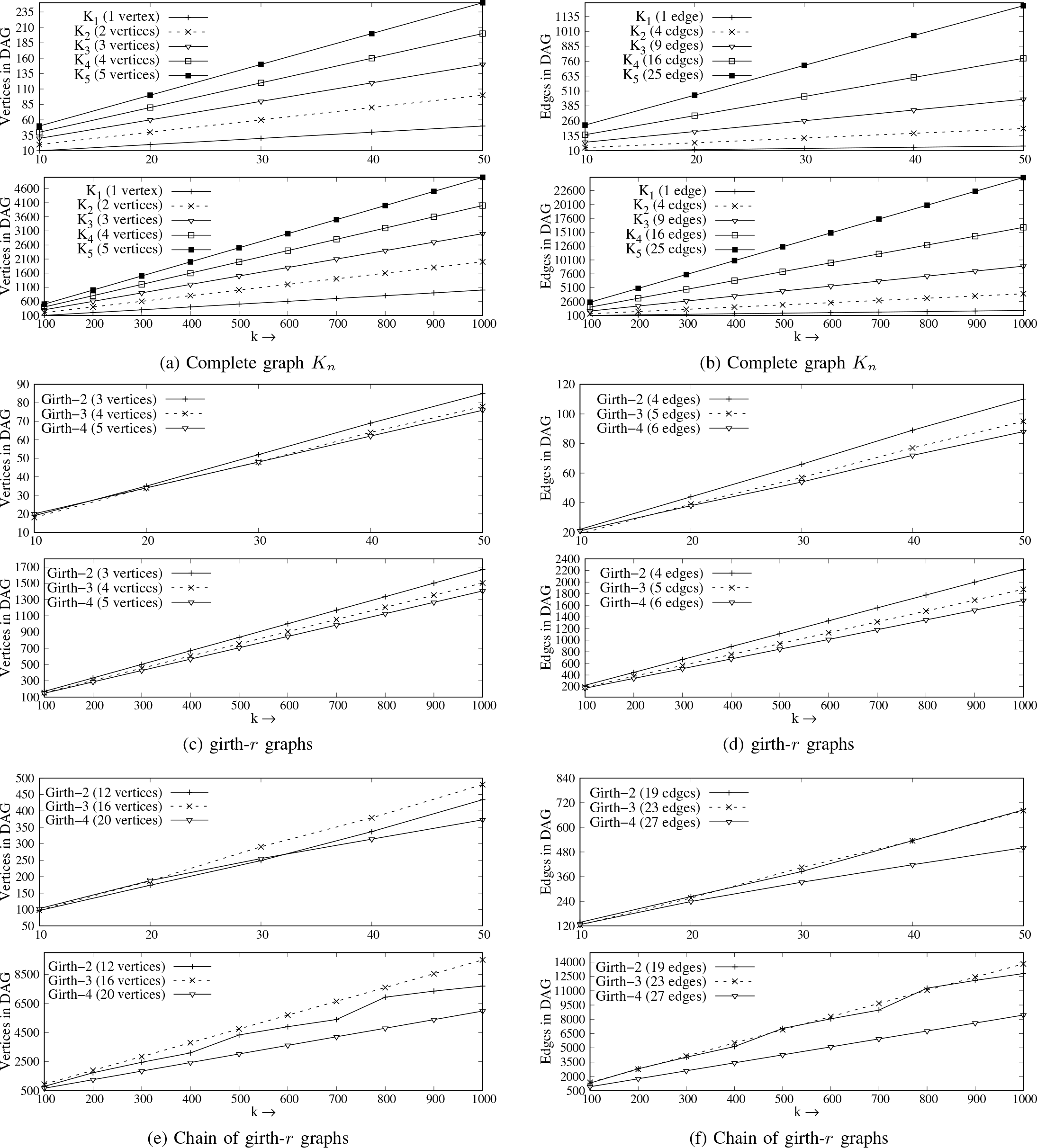
Increase in vertices and edges due to DAGification for the graphs in 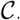.

The increase in edges and vertices are separately plotted in Figure 3. As seen from these plots, both edge and vertex cardinalities increase linearly with *k*. The increase is lesser for graphs with larger girth. For example, Figures 3e and 3f shows that though the girth 4 graph has 27 edges and girth 2 graph has 19 edges, the DAG output for the latter has significantly higher size than the former for every *k* value.

From these plots we see that the DAGification produces graphs with much larger size than the original input graph. This affects both the preprocessing cost and the subsequent alignment cost for POA based alignment. Recall that the worst case performance of V-ALIGN, even with the naive feedback vertex set computation, is *O*(*kVE*) where *k* is the sequence length and *V* and *E* are the vertex and edge counts of the input graph. On the other hand, the POA alignment cost is *O*(*kE*′) where *E*′ is the edge count of the DAG. From the plots, we see that in *E*′ is significantly more than *V* · *E* in most cases. For example, the chain of girth 2 graphs has *VE* = 228 where as *E*′ = 13, 814 (DAG size in Figure 3f for *k* = 1000). Similarly, *VE* = 12 for girth 2 graph and *E*′ = 1877 (DAG size in Figure 3d for *k* = 1000). For the complete graph *K*_5_, *VE* = 125 and *E*′ = 24, 975 (DAG size for *k* = 1000). That is, already for sequence length *k* = 1000, the computational steps for POA based alignments increases by roughly 100 fold or more as compared to V-ALIGN in several graphs.

## VI. CONCLUSION

We give a novel alignment algorithm V-ALIGN that directly aligns an input sequence onto a directed graph without requiring DAGification. V-ALIGN is based on a novel dynamic programming formulation that allows gapped alignment (constant, linear or affine) directly on the input graph. The time to fill the DP table has linear dependence on the sizes of the sequence, the graph and its feedback vertex set. Our experiments show that V-ALIGN achieves considerable saving in the computational steps compared to POA based alignment. The saving is even more significant for larger input sequence lengths. When the alignment is restricted to a filtered set of small subgraphs for improved efficiency, the V-ALIGN can be used for aligning to these candidate subgraphs. In this case, its performance depends on the subgraph sizes.

